# Protein Kinase C and the stress response pathways are required for kinetochore homeostasis

**DOI:** 10.1101/078196

**Authors:** Elena Ledesma-Fernández, Eva Herrero, Guðjón Ólafsson, Peter H Thorpe

## Abstract

Kinetochores serve both a structural role linking chromosomes to the mitotic spindle and a regulatory role, controlling the timing of mitosis via the spindle assembly checkpoint. To identify proteins that regulate the kinetochore we used a genome-wide fluorescence microscopy approach. We combined an array of mutants of either non-essential gene deletions or essential temperature-sensitive alleles with fluorescently tagged spindle pole bodies (centrosome) and outer kinetochores. Quantitative and qualitative analysis revealed mutants that affect the levels and distribution of kinetochores respectively. These mutants are enriched for those involved in mRNA processing, chromatin organization, DNA replication/repair and mitosis. Our data show that the Pkc1 kinase maintains the kinetochore focus via its ability to prevent cell stress and this phenotype is rescued by an osmotic stabilizer. These data support the notion that kinetochore and microtubule homeostasis are perturbed by the stress response pathways. Hence this observation provides a candidate mechanism for extracellular stress leading to chromosome segregation defects.

## Introduction

Kinetochores are large mega-dalton complexes that assemble on chromosomes at the centromeres and direct chromosome segregation (Biggins, 2013; Cheeseman, 2014). Defects in chromosome segregation lead to aneuploidy, in which daughter cells have gained and/or lost whole chromosomes. Aneuploid cells are a hallmark of cancer tissue and aneuploidy may aid aspects of tumourigenesis or tumour evolution (de Bruin et al, 2014; Sotillo et al, 2007; Weaver et al, 2007). To prevent aneuploidy, cells carefully regulate the structure and function of their kinetochores to ensure accurate chromosome segregation. The budding yeast kinetochore provides perhaps the simplest model of a eukaryotic kinetochore, since it assembles on a single nucleosome of a short ‘point’ centromere, yet many of the genes encoding yeast kinetochore proteins are well conserved with those of mammals. Despite the simplicity of the budding yeast kinetochore, it is a large complex, consisting of at least 60 different proteins, many present in multiple copies (Biggins, 2013; Westermann et al, 2007). The first step in assembling kinetochores is the loading of a centromere specific variant of histone H3, Cse4, which then recruits the remaining kinetochore proteins (Collins et al, 2005). The key function of the fully assembled kinetochore is to link chromosomes to microtubules (MTs); this function is achieved primarily by the NDC80 complex (Cheeseman et al, 2006; DeLuca & Musacchio, 2012). The kinetochore’s ability to maintain MT attachment is complicated by the dynamically extending and shrinking nature of the ‘plus’ end of the MT. The heterodecameric DAM1-DASH complex is critical for this function and forms the most distal part of the kinetochore from the centromeric nucleosome (Joglekar et al, 2006; Shang et al, 2003). The DAM1-DASH complex requires the NDC80 complex for loading onto microtubules (Lampert et al, 2013; Maure et al, 2011). Sixteen DAM1-DASH decamers can encircle a MT *in vitro*, forming a ring (Gonen et al, 2012; Miranda et al, 2005; Ramey et al, 2011; Wang et al, 2007; Westermann et al, 2005). The genes encoding components of the DAM1-DASH complex are not conserved in mammalian cells, but the SKA complex appear to serve an analogous function, for example see (Jeyaprakash et al, 2012).

Although the constituents of the kinetochore are well characterized (Biggins, 2013; Westermann et al, 2007), the proteins that regulate its function are less clearly defined. Key exceptions are the spindle assembly checkpoint (SAC) and the Aurora kinase B, which act by phosphorylating specific kinetochore proteins to delay anaphase or release microtubules respectively, for reviews see (Carmena et al, 2012; Jia et al, 2013; Musacchio & Salmon, 2007). There are multiple other examples of protein modifications affecting kinetochore function, for example methylation by Set1 (Zhang et al, 2005), ubiquitylation by Psh1 (Hewawasam et al, 2010; Ranjitkar et al, 2010) or phosphorylation by the ATR orthologue, Mec1 (Strecker et al, 2016). However, we predict that other proteins that regulate kinetochores remain to be identified, based upon the large groups of proteins that have been found to affect the morphology of the mitotic spindle (Vizeacoumar et al, 2010) and induce a chromosomal instability (CIN) phenotype (Stirling et al, 2011). In an attempt to identify novel regulators of the kinetochore, we performed a genome-wide screen of mutants of both non-essential and essential genes to detect changes in the levels and localization of the DAM1-DASH component, Dad4. Our analysis reveals a set of mutants that are enriched for those involved in chromosome segregation but also, DNA replication and repair, chromatin organization, mRNA processing and splicing. We were surprised to find a Dad4-YFP phenotype produced both by mutants that affect actin and the cell wall integrity pathway kinase, Pkc1. *PKC1* mutants produce a CIN phenotype (Stirling et al, 2011), therefore we examined their kinetochore phenotype in more detail. We find that *PKC1’s* effect upon the kinetochore likely acts via actin and MT stability, since the phenotype can be recapitulated with mutants or drugs that disrupt their function. This work establishes a mechanistic link between the stress response pathways and the kinetochore/mitotic spindle and suggests that stress plays an important role in kinetochore homeostasis.

## Results

### A screen for aberrant kinetochores

To identify mutants that affect kinetochore function we created yeast strains encoding Dad4 fused to yellow fluorescent protein (Dad4-YFP) and Spc42 fused with red fluorescent protein (Spc42-RFP). Dad4 is a member of the outer kinetochore DAM1-DASH complex and Spc42 is a structural protein within the spindle pole body (SPB). Both of these fluorescently-tagged alleles are at their endogenous loci under the control of their native promoters. Fluorescence imaging shows stereotypical yeast kinetochore and SPB foci (Fig 1A). These yeast strains also contain haploid specific marker genes *(Tong & Boone, 2007)* to allow us to use the *Synthetic Genetic Array* (SGA) methodology (Tong et al, 2001) to combine the two fluorescently-tagged proteins with arrays of mutants.

**Figure 1.**
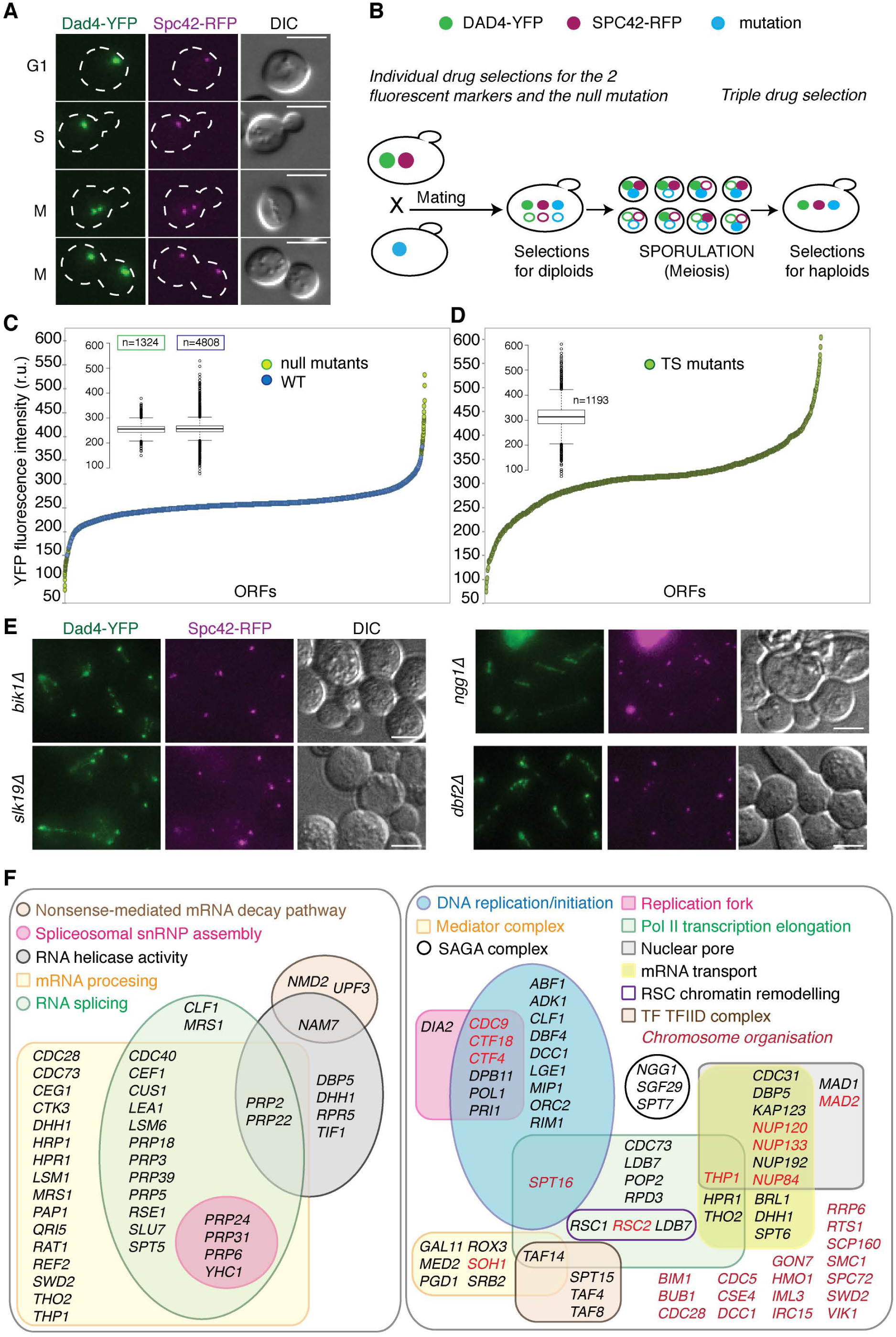
A genome-wide screen for mutants that disrupt the kinetochore. A Endogenously-encoded Dad4-YFP and Spc42-RFP allow us to visualize kinetochores and spindle pole bodies using fluorescence imaging at different stages of the cell cycle; the scale bars are 5 µm. B The strategy for Synthetic Genetic Array (SGA) analysis is illustrated. The haploid yeast strain encoding Dad4-YFP (green circle) and Spc42-RFP (purple circle) is mated to an array of strains each of which contain a specific mutation (blue circle). The resulting diploids are sporulated and the haploid progeny containing both a mutation and two tagged genes is selected. C The plot shows the quantitation of the levels of Dad4-YFP in the non-essential gene deletions from the lowest intensity (left) to the highest intensity (right). Each point represents the mean fluorescent intensity of kinetochore foci for each strain. Control (WT) strains are indicated in blue and mutants in green. The box-and-whisker plot (inset) shows the distribution of the data of both the controls and mutants. The box and whiskers plots indicate the median (bar), quartiles (box) and 1.5 times the interquartile range (whiskers), with outliers plotted as dots. D The plot shows the quantitation of the levels of Dad4-YFP for the temperature-sensitive (ts) strains plotted from the lowest intensity (left) to the highest intensity (right). The box and whiskers plot (inset) shows the distribution of the fluorescence for the ts mutants, as for panel C. E Four examples of mutants that produce a ‘visible’ Dad4-YFP phenotype are shown. *bik1∆*, *slk19∆*, *ngg1∆* and *dbf2∆* strains all have abnormal distribution of Dad4-YFP. The scale bars are 5 µm. F Gene Ontology enrichment analysis of the mutants that perturb Dad4-YFP levels or localization reveal categories of proteins that we had not anticipated to be related to kinetochore function. These included m*RNA processing* (*p*=3×10^−13^), *RNA splicing* (*p*=6×10^−7^) and *spliceosomal snRNP assembly* (*p*=5×10^−4^) (left panel). The mutants were also enriched for those involved in the *mitotic cell cycle* (*p*=4×10^−14^); including *DNA replication* (*p*=1×10^−4^), PolII transcription elongation (*p*=7×10^−5^) and the *nuclear pore* (*p*=1×10^−4^) (right panel). Source data for this figure is available on the online supplementary information page.

We used SGA to combine the *DAD4-YFP* and *SPC42-RFP* alleles with the non-essential deletion collection (Winzeler et al, 1999), which includes 4746 separate viable deletion strains, to systematically assess the effect of specific mutations on the kinetochore (Fig 1B). The resulting haploid strains were imaged on agar pads using wide-field epi-flourescence microscopy. We included 1302 copies of a control strain to assess variability of the assay (a deletion of the *HIS3* gene, which is also absent in the BY4742 genetic background of the other deletions strains). To examine the impact of essential genes on the kinetochore, we employed a collection of temperature-sensitive (ts) alleles (Li et al, 2011) (1334 members, include 754 unique alleles of 475 unique ORFs). These strains were also combined with the fluorescence alleles using SGA and the resulting haploid strains were then grown for 5 hours at 37°C prior to imaging. Since we used a high numerical aperture objective lens (63x, 1.4NA), we were able to optically section through the yeast cells to produce 3-dimensional images (all of the image data from this screen is available via the Open Microscopy Environment, Image Data Repository, idr-demo.openmicroscopy.org with the accession identifier ‘idr0011’). These images were then analyzed using a fully automated image analysis protocol (Ledesma-Fernandez & Thorpe, 2015); we quantified the fluorescence intensity of each focus in each image. The fluorescence intensity is a measure of the local protein concentration at the kinetochore and has been widely used to measure kinetochore protein abundance (Joglekar et al, 2008; Joglekar et al, 2006).

The kinetochore intensities for all the strains analyzed shows a near-normal distribution and the yeast strains that differ most from the overall mean are enriched for mutants over controls (Fig 1C and Fig 1 source data). For the essential ts mutants we found a larger spread of Dad4-YFP fluorescence (Fig 1D). In addition to the fully-automated quantitative analysis, we also manually examined all of the images to determine which did not have a standard stereotyped kinetochore localization of Dad4-YFP. Such a kinetochore phenotype would potentially be missed by quantitative measurements. We found 85 non-essential mutants that had such a phenotype (for examples see Fig 1E and Fig 1 source data). Many of these (48 out of the 85) were already identified in our quantitative screen, but 37 were missed with quantitative analysis. We did not include any of the essential ts mutants in this “visual foci phenotype” category since most strains that we examined would have scored as having aberrant kinetochores.

We retested 338 non-essential mutant strains that included the most extreme variants in Dad4-YFP intensity (high and low) and strains with a visual kinetochore phenotype, 45 copies of the control strain and 146 strains that we chose to retest based upon either their annotations related to spindle function or established links with the other high/low-Dad4-YFP mutants. Of 224 non-essential mutants that produced a Dad4-YFP phenotype in the primary screen (high-intensity, low-intensity or visual) and that we were able to score in the secondary screen, 192 consistently gave a phenotype (86%, Fig 1 source data). Additionally, we were unable to score 20 mutants in the secondary screen that produced a Dad4-YFP phenotype in the primary screen and, based upon the high validation rate, we include these in our final list of mutants affecting Dad4-YFP. Consequently, our screen identifies 175 non-essential deletion mutants that affect Dad4-YFP levels (72 that decrease Dad4-YFP foci intensity, 103 that increase intensity; Fig 1 source data) and 37 additional mutants that produce a visual phenotype. We retested essential temperature sensitive alleles that had Dad4-YFP levels <200 relative units and > 400 relative units. Strains that had aberrant Dad4-YFP levels in both the primary and retest screens were considered further (Fig 1 source data). 67 essential alleles decrease Dad4-YFP foci intensity and 48 increase it. Interestingly, some of the alleles of the ts collection are present in multiple copies since they were acquired from different sources. We find that these alleles can sometimes differ with one giving high intensity Dad4-YFP foci and the other giving low intensity foci (referred to as “low, high” in Fig 1 source data). An example of this is the *cdc20-1* allele, which consistently gave both high and low intensity foci from different isolates. In total 116 ts alleles gave a Dad4-YFP intensity phenotype, corresponding to 102 separate open reading frames. We found that a large proportion (~27%) of the essential alleles affect Dad4-YFP levels (Fig 1D and Fig 1 source data), which contrasts with only ~3% of non-essential genes affecting Dad4-YFP levels. In total, therefore, we have identified 314 genes, whose mutants produce a Dad4-YFP phenotype (Fig 1 source data).

## Global analysis

To examine these mutants collectively we asked which gene ontology terms are enriched using the GOrilla algorithm (http://cbl-gorilla.cs.technion.ac.il/ (Eden et al, 2009)). This analysis revealed enrichment of categories predicted to impact kinetochore function such as *chromosome segregation* and *regulation of chromosome segregation* (enrichment *p* values 4×10^−5^ and 5×10^−7^, respectively) We also found that enriched categories include terms that we had not anticipated Fig 1F e.g. *mRNA Processing*, *RNA helicase activity and splicing.* These include three key members of the nonsense-mediated mRNA decay (NMD) pathway (*NMD2*, *NAM7* and *UPF3*). In addition, we found enrichment categories involved in *chromatin organization, DNA replication, transcription elongation* and the *nuclear pore* (Fig 1F). For example, we found six members of the mediator complex (*MED2*, *PGD1*, *ROX3*, *SRB2, SOH1* and *TAF14*), three members of the RSC chromatin-remodeling complex (*RSC1*, *RSC2* and *LDB7*) and three members of the SAGA complex (*NGG1*, *SGF29* and *SPT7*).

We expected CIN mutants would show significant variability in their kinetochore foci intensities from cell to cell since we have previously found this to be true in selected cases where the intensity of Dad4-YFP was increased, for example *hmo1∆* and *mad1∆* (Berry et al, 2016; Ledesma-Fernandez & Thorpe, 2015) both of which were identified in this screen. Consistent with this notion, we identified 98 of the 723 CIN genes (Stirling et al, 2011), an enrichment *p*-value of 10^−14^. We also note that a number of mutants were identified in the threonine/serine biosynthetic pathway: *hom2∆*, *hom3∆, aat2∆ and thr1∆* (Fig S1 A, Fig 1 source data). For example, *hom2∆* cells showed increased and highly variable levels or Dad-YFP at kinetochore focus (Fig S1 B, C and D). However, our media for both the SGA and microscopy does not contain serine or threonine. Supplementing the media with those amino acids rescued the kinetochore phenotype (Fig S1 B, C and D). Other notable complexes identified in this Dad4-YFP screen were members of the RSC and SWI/SNF chromatin remodeling complexes (Fig S2), the mitotic exit network (Fig 1E *dbf2∆* and Fig 1 source data), microtubule associated proteins and the kinetochore (Fig S3). Mutants of kinetochore and associated genes gave diverse Dad4-YFP phenotypes, for example *kre28∆*, *ctf19∆*, *bim1∆*, *bik1∆* and *slk19∆* produced a Dad4-YFP signal stretching between the spindle poles (Fig 1E and Fig S3) consistent with microtubule localization. In contrast *ask1-2* and *spc34-ts* have elevated levels of Dad4-YFP in their kinetochore foci, whereas *dam1-5* and *duo1-2* reduce the levels of Dad4-YFP (Fig S3F). We also identified multiple components of the *Nonsense-Mediated mRNA decay* pathway (NMD) that serves to degrade RNA and regulate gene expression (He et al 1997). Several kinetochore RNAs have been shown to increase in NMD mutants (Dahlseid et al 19898, Kebaara et al 2009). Deletion of the NMD genes *UPF3*, *NAM7* and *NMD2* all had high levels of Dad4-YFP with considerable variability (Fig S4).

## Protein Kinase C

One of the 107 CIN mutants that affected Dad4-YFP was an allele of the *PKC1* gene, which encodes the only Protein Kinase C in budding yeast (Pkc1). Pkc1 is an essential kinase that functions in the Cell Wall Integrity (CWI) pathway in addition to multiple other roles throughout the cell (Levin, 2005; Levin et al, 1990). Mammalian cells have multiple PKC isoforms, which also have diverse functions (Mellor & Parker, 1998). Mutations in yeast *PKC1* have a chromosomal instability phenotype (Stirling et al, 2011) despite Pkc1 localizing to the cell periphery and bud neck. Three *PKC1* temperature-sensitive mutant alleles were tested in the screen and we found the more severe the phenotype the lower the reported restrictive temperature for the allele (Fig 1 source data). We retested the *pkc1-1* strain after 5 hours growth at either 23°C or 37°C. At the higher temperature we confirmed that many of the *pkc1-1* cells had lost their Dad4-YFP foci whereas Spc42-RFP foci was still present (Fig 2A and B). To confirm that the Dad4-YFP phenotype in *pkc1-1* cells was caused by a failure in Pkc1, we next added back a wild-type copy of the *PKC1* gene at the *URA3* locus and found that this was sufficient to rescue the Dad4-YFP foci phenotype in the *pkc1-1* strain at 37°C (Fig 2C and D).

**Figure 2.**
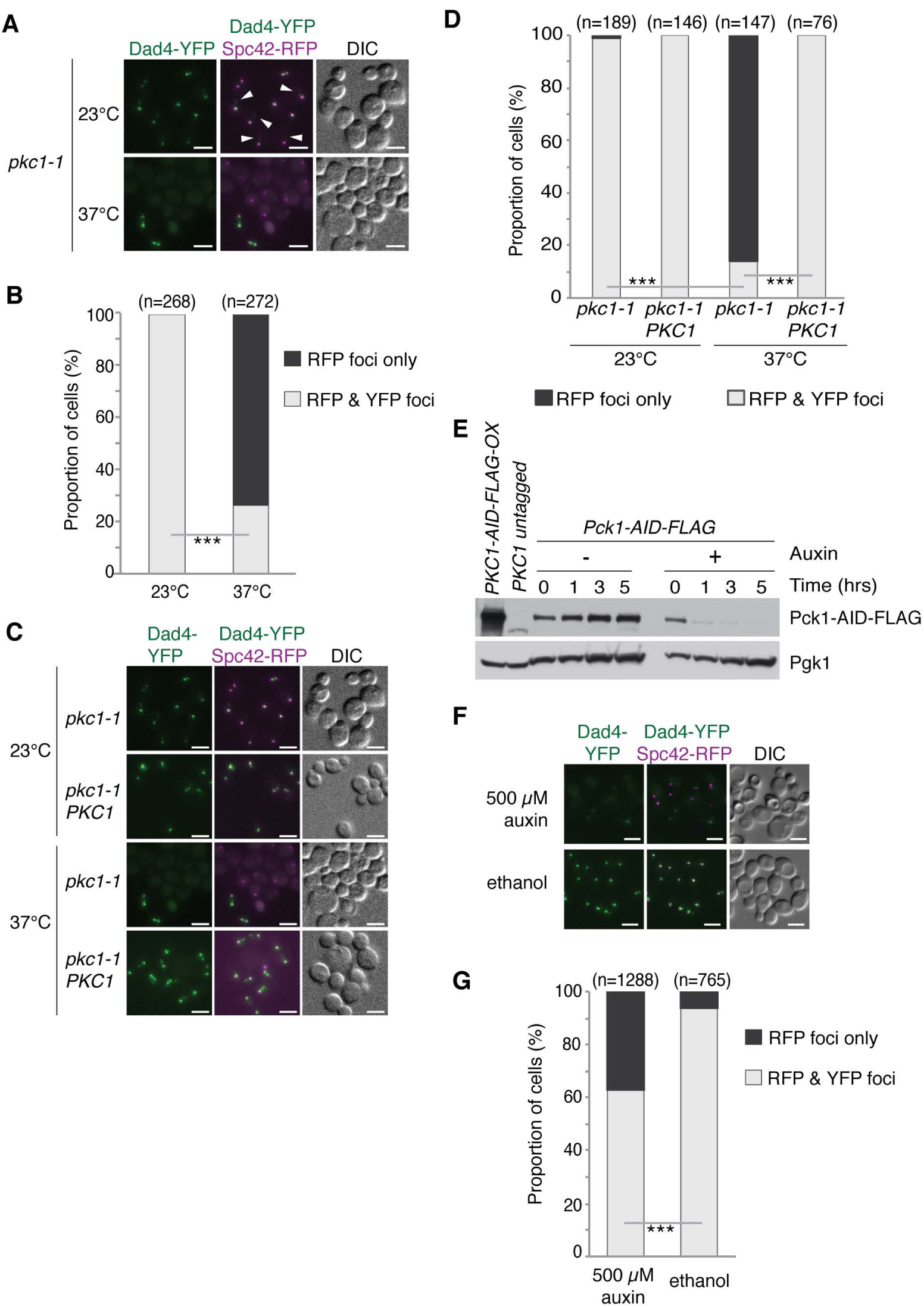
Disruption of *PKC1* affects the level of Dad4-YFP. A Dad4-YFP distribution in the *pkc1-1* strain. At 23°C, Dad4-YFP kinetochore foci are largely normal, except for a few additional weak foci in some cells (white arrowheads). Upon shifting to 37°C for 5 hours most *pkc1-1* cells have lost their Dad4-YFP foci. Scale bars are 5 µm. B The proportion of *pkc1-1* cells that have lost their Dad4-YFP foci is plotted after growth for 5 hours at either 23°C or 37°C. *** indicates a *p* value of 3×10^−25^ from a Fishers exact test. C The kinetochore phenotype caused by disrupting Pkc1 with the *pkc1-1* mutation at 37 °C is mitigated by the introduction of a wild-type copy of the *PKC1* gene at the *URA3* locus. Scale bars are 5 µm. D The proportion of *pkc1-1* and complemented *pkc1-1 PKC1* cells that have lost their Dad4-YFP foci is plotted after growth for 5 hours at either 23°C or 37°C. *** indicates *p* values of less than 10^−40^ from Fisher exact tests. E Analysis of whole cell protein extracts show that Pkc1-AID protein is depleted from cells after addition of 500 µM auxin. Phosphoglycerate kinase (Pgk1) is used as a loading control. Whole cell extract from cells overexpressing *PKC1-AID-FLAG* (first lane) and from cells with untagged *PKC1* (second lane) were used as positive and negative control, respectively. F The Dad4-YFP phenotype of *pkc1-1* cells can be largely recapitulated by depleting Pkc1 after addition of auxin. Scale bars are 5 µm. G The proportion of PKC1-AID cells that have lost their Dad4-YFP foci after 3 hours of treatment with 500 µM auxin. *** indicates a *p* value of less than 10^−50^ from a Fishers exact test. We note that there may be some Pkc1-AID degradation without auxin, since some cells lose Dad4-YFP foci in the ethanol control.

To confirm that the kinetochore phenotype seen in *pkc1-1* cells was not dependent upon high-temperature, we used an ‘auxin-induced degradation’ (AID) system (Morawska & Ulrich, 2013; Nishimura et al, 2009). This approach uses a short AID tag fused to Pkc1 and AID-specific E3 ubiquitin ligase (AFB2). The interaction between the AID domain and AFB2 is dependent upon the plant hormone auxin (indole-3-acetic acid). In a strain encoding Dad4-YFP and Spc42-RFP, we fused the endogenous *PKC1* gene with an AID tag (*PKC1-AID*) and integrated the *AFB2* gene at the *URA3* locus. We found that after addition of 500 µM auxin, Pkc1 is depleted (Fig 2E) and the proportion of cells without visible Dad4-YFP foci increased (Fig 2F and G). These data confirm that the kinetochore phenotype of the *pkc1-1* can be recapitulated by depletion of Pkc1. We noted that the *AFB2* gene was particularly unstable in this strain suggesting that Pkc1 degradation is partially active even without auxin, a previously reported effect (Morawska & Ulrich, 2013). This notion is consistent with the small loss of Dad4-YFP in cells without auxin (Fig 2G), but we note that the protein analysis suggests that most cells retain Pkc1 (Fig 2E).

A potential mechanism to explain the *pkc1-1* kinetochore phenotype is reduced expression of kinetochore genes or degradation of kinetochore proteins.

However, we confirmed that several kinetochore proteins (Dad4, Dad3 and Mtw1) are not degraded in *pkc1-1* cells at the restrictive temperature (Fig S5A)

Since disruption of Pkc1 has an effect on the Dad4 protein levels at the kinetochore, we asked whether this effect extended to other members of the DAM1-DASH complex and kinetochore or was specific to Dad4. We transferred the *pkc1-1::KANMX* allele into strains encoding Dad3-YFP, Mtw1-YFP or Ndc10-GFP representing kinetochore components in the outer DAM1-DASH, central and inner kinetochore complexes respectively (Fig 3A). Upon shifting to the higher temperature we found that, in each case, the proportion of cells with visible kinetochore foci decreased (Fig 3B, C and D). Thus, the effect of perturbing Pkc1 is not specific to Dad4 or the DAM1-DASH complex, but extends to the inner kinetochore proteins. Thus, mutation of *PKC1* is sufficient to cause an aberrant kinetochore phenotype.

**Figure 3.**
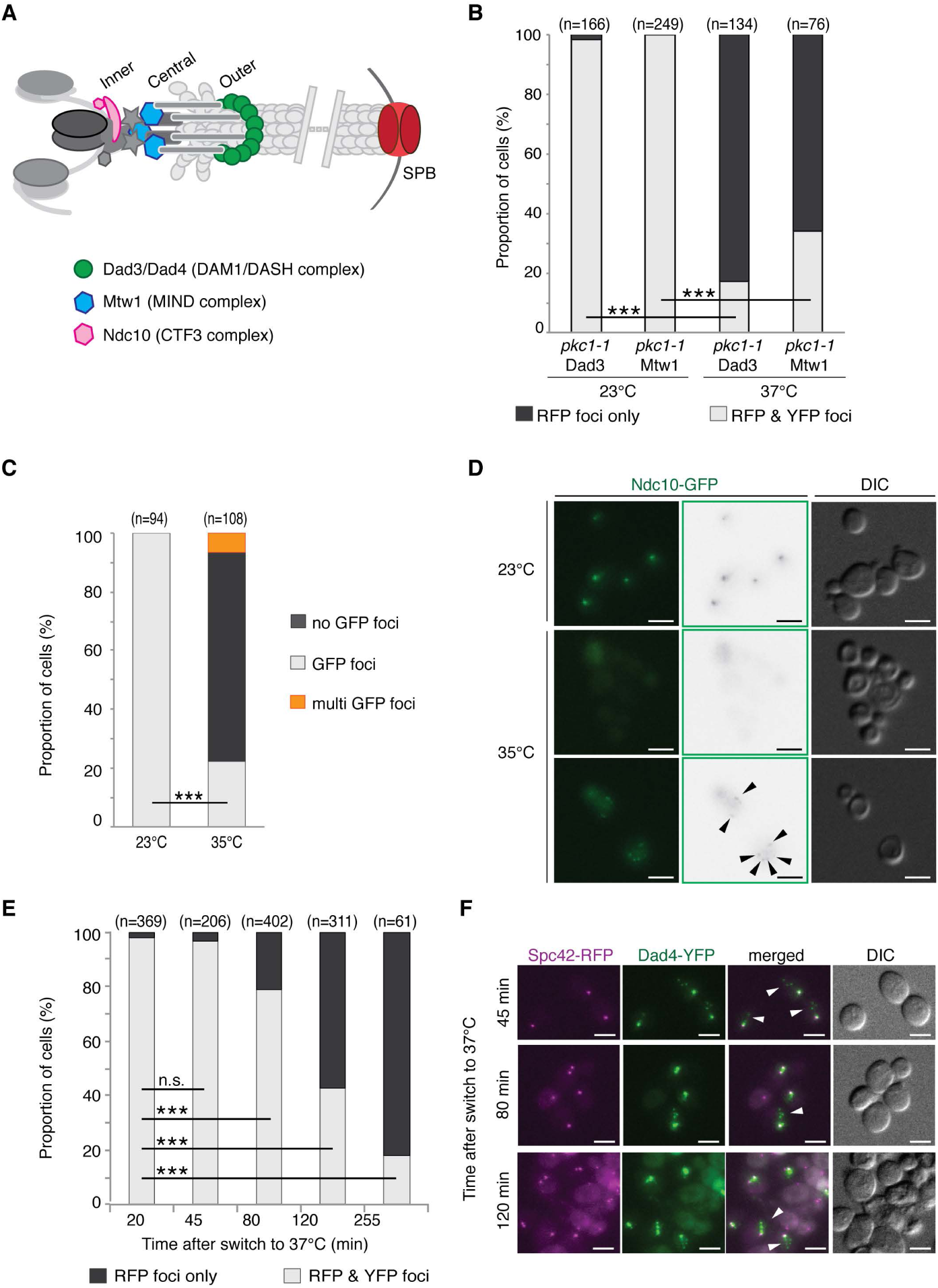
Disruption of *PKC1* leads to kinetochore declustering. A The schematic illustrates three additional fluorescently-tagged kinetochore proteins that were analyzed in *pkc1-1* cells. Ndc10 is a member of the CBF3 inner kinetochore complex, Mtw1 is part of the central MIND complex and Dad3, like Dad4, is a member of the DAM1-DASH complex at the outer kinetochore. B The proportions of *pkc1-1* cells that have lost Dad3-YFP or Mtw1-YFP kinetochore foci are plotted after growth for 5 hours at either 23°C or 37°C. *** indicates *p* values of less than 10^−40^ from Fishers exact test. C The proportions of *pkc1-1* cells that have lost their Ndc10-GFP foci are plotted after growth for 5 hours at either 23°C or 35°C. We note that in ~5% of the *pkc1-1* cells multiple Ndc10-GFP foci were observed at 35°C (orange category). *** indicates a *p* value of 1×10^−30^ from a Fishers exact test. D Examples of *NDC10-GFP pkc1-1* cells at 23°C and 35°C are shown. In the lower panels we include a cell that shows the ‘multi-foci’ phenotype of Ndc10-GFP (black arrowheads); the scale bars are 5 µm. E The proportions of *pkc1-1* cells that have lost their Dad4-YFP foci after growth for varying times at 37°C are plotted. n.s. indicates not statistically significant and *** indicates *p* values of less than 10^−17^ all using Fishers exact test. F Prior to complete loss of the Dad4-YFP, we observed *pkc1-1* cells with weak extra Dad4-YFP foci (white arrowheads); the scale bars are 5 µm, showing that the kinetochore foci subdivide after switching to the restrictive temperature.

We noticed that in *pkc1-1* cells, small additional Dad4-YFP foci could be observed even at the lower temperature (Fig 2A and C). Also Dad4-YFP foci in the *pkc1-1 ura3-1::PKC1::URA3* strain do not appear entirely normal, with multiple foci in some cells (Fig 2C). This suggests that non-lethal disruption of Pkc1 is sufficient to perturb the kinetochore. In yeast all 16 centromeres/kinetochores cluster together to produce the distinctive kinetochore focus. The additional kinetochore foci lead us to ask whether the *pkc1-1* kinetochore phenotype is consistent with kinetochore de-clustering. To test this notion we monitored the Dad4-YFP phenotype over time after shifting *pkc1-1* cells to the non-permissive temperature. We find that the proportion of cells with Dad4-YFP foci decreases over time (Fig 3E). Furthermore, small disperse foci appear in cells at early time points, consistent with de-clustering of kinetochores after the shift to the non-permissive temperature (Fig 3F).

*PKC1* is an essential gene whose product functions downstream of Rho1 in the CWI pathway (Fig 4A) (Levin et al, 1990). *PKC1* mutants die due to osmotic stress and this lethality can be rescued by the addition of an osmotic stabilizer, such as 1 M sorbitol (Paravicini et al, 1992). To test whether the kinetochore phenotype was linked to a failure in CWI, we attempted to rescue the kinetochore phenotype of *pkc1-1* cells with sorbitol. *pkc1-1* cells were grown at 37°C for 5 hours with or without sorbitol and Dad4-YFP signal assessed by microscopy. We found that 1 M sorbitol is sufficient to rescue the visible disappearance of Dad4 foci (Fig 4B and C), however, the rescued cells had additional foci suggesting either non-kinetochore clusters of Dad4 or partially declustered centromeres.

**Figure 4.**
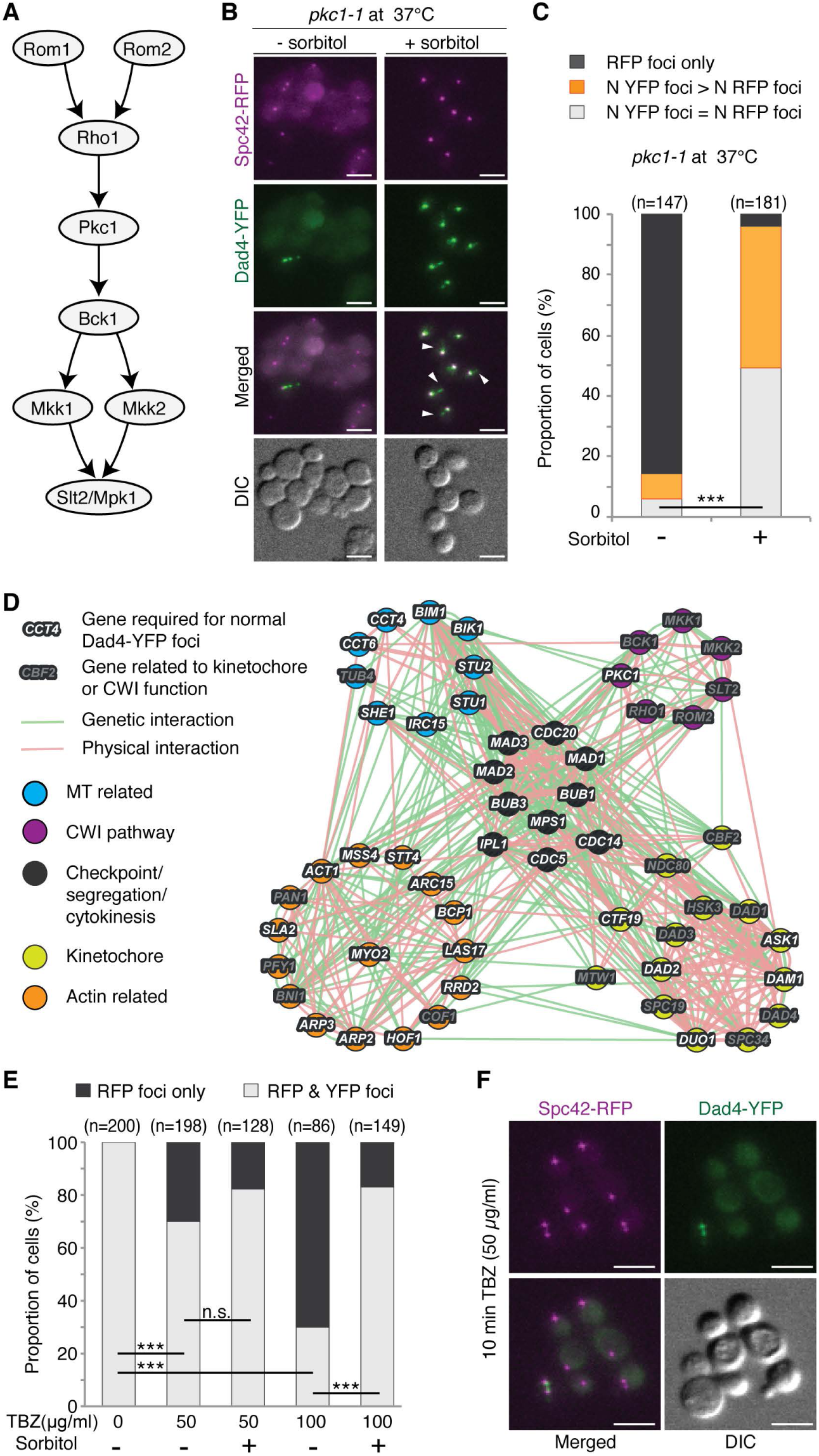
Cell stress affects kinetochore protein levels. A Pkc1 is part of a MAP-kinase signaling cascade of the Cell Wall Integrity (CWI) pathway. Pkc1 is activated by Rho1 and acts directly upon Bck1. B The Dad4-YFP phenotype observed in *pkc1-1* cells, after 5 hours of growth at 37°C, is largely rescued by addition of 1M sorbitol. Although, we noted that many *pkc1-1* cells in sorbitol had multiple Dad4-YFP foci (white arrowheads); scale bars are 5 µm. C The proportion of *pkc1-1* cells that have lost their Dad4-YFP foci are plotted after growth for varying times at 37°C with and without sorbitol. Nearly half of the *pkc1-1* cells in sorbitol have multiple Dad4-YFP foci. *** indicates a *p* value of 4×10^−19^ using Fishers exact test. D Actin and microtubule (MT) mutants also cause a Dad4-YFP kinetochore phenotype (Fig 1 source data). The nodes of the interaction map depict genes whose mutants we identified to affect Dad4 levels or localization (light text) or genes not identified in the screen but related to CWI or kinetochore function (dark text). Genes are grouped and colour-coded based upon their function. Genes are linked by genetic (green lines) and physical (red lines) interactions. E The proportions of wild type (WT) cells that have lost their Dad4-YFP foci are plotted after growth with and without the microtubule-destabilizing drug thiabendazole (TBZ) and sorbitol. TBZ causes a proportion of cells to lose their Dad4 foci. At high concentration (100 µg/ml) TBZ produces a strong kinetochore phenotype that can be partially rescued with sorbitol. n.s. indicates not statistically significant (*p*=0.076) and *** indicates *p* values less than 10^−15^ all using Fishers exact test. F An example of WT cells treated with 50 µg/ml TBZ for 10 minutes is shown. The scale bars are 5 µm.

These data suggest that osmotic stress contributes significantly to the Dad4-YFP phenotype. However, mutants of the other members of the CWI pathway (Rom2, Bck1, Mkk1, Mkk2 and Slt2) show no comparable Dad4-YFP phenotype with the exception of *stl2Δ* (Fig S5B). We asked whether the canonical stress response pathway (Hog1, p38) is active in *pkc1-1* cells. Hog1 is the canonical mitogen-activated protein (MAP) kinase in yeast and is activated in response to stress (Brewster et al, 1993). We probed whole cell extracts from *pkc1-1* cells and find that Hog1 is phosphorylated consistent with an activated stress response pathway although both at the permissive and restrictive temperatures (Fig S5C). These data show that while the Dad4 phenotype of *pkc1-1* mutants is linked to stress, this is not entirely dependent upon the CWI MAP kinase cascade pathway.

Since sorbitol was able to partially rescue the Dad4 phenotype of *PKC1* mutants we looked for a connection between osmotic/cell wall stress and spindle biology. We noted that in the Dad4-YFP screen we identified a class of mutants related to actin dynamics (Fig 4D, Fig 1 source data). These include actin itself (e.g. *act1-101/105/121/133/136/159*), myosin (*myo2-16*), members of the Arp2/3 complex (e.g. *arp2-14*, *arp3-F306G*, *las17-13*) and others (Fig 1 source data). There is precedent for cytoskeletal defects leading to MT defects in fission yeast, where osmotic stress induces depolymerisation of the actin cytoskeleton and a perturbation in microtubule dynamics (Robertson & Hagan, 2008). Furthermore, mutants that affect the cell wall are important for MT stability (*LAS17*, *ECM27*, *SUR7*, *ECM33*, *ECM8*, *HLR1* and *KTR1*) (Vizeacoumar et al, 2010). If both *pkc1-1* and the actin mutants disrupt normal microtubule dynamics then we expected that a direct disruption of microtubule function would lead to a Dad4 phenotype.

Indeed, we identified a number of MT associated proteins required for normal Dad4-YFP levels or localization: Bim1, Bik1, Stu2, She1, Irc15, Rrd2 (Fig 1 source data). To test this notion we used the microtubule destabilizing drug thiabendazole (TBZ) to examine the effects upon Dad4-YFP foci. After addition of increasing amounts of TBZ we saw an increase in cells lacking Dad4-YFP foci (Fig 4E and F). Interestingly, we noted that this phenotype is partially rescued by the addition of 1 M sorbitol, suggesting that sorbitol may do more than simply protect cells from osmotic shock (Fig 4E). These data highlight two points, first that disruption of MTs or MT dynamics could underlie Dad4 kinetochore phenotypes in the *pkc1-1* and actin mutants, and second that sorbitol may stabilize MTs – a notion supported by previous studies (Gerson-Gurwitz et al, 2009; Korolyev et al, 2005).

## Synthetic Physical Interactions

There is evidence that Pkc1 may act on nuclear proteins directly, since removal of a domain from the N-terminus of Pkc1 (HR1) leads to its localization to the mitotic spindle (Denis & Cyert, 2005). Furthermore, there are reported physical interactions between Pkc1 and the kinetochore proteins Nnf1, Nuf2 and Spc105 (Wang et al, 2012). We did not detect Pkc1 at the kinetochore by fluorescence imaging of a Pkc1-GFP strain (Fig S5D) and Pkc1 has not been isolated with purified kinetochores (Akiyoshi et al, 2010; Ranjitkar et al, 2010). However, we wanted to ask whether forced recruitment of Pkc1 to the kinetochore would lead to a mitotic phenotype. To test this, we made use of the *Synthetic Physical Interaction* (SPI) system to separately associate Pkc1 with kinetochore and associated proteins (Olafsson & Thorpe, 2015). The SPI system uses an antibody fragment that binds to GFP, the GFP-binding protein (GBP), which is fused to Pkc1, to bind to GFP-tagged proteins in the cell. The GBP includes an RFP tag to enable colocalization to be monitored by fluorescence imaging. We used plasmids encoding the GBP alone, Pkc1-GBP (wildtype Pkc1), *pkc1-ha*-GBP (a constitutively hyperactive mutant of Pkc1, *pkc1-R398A-R405A-K406A* (Martin et al, 2000) and *pkc1-kd*-GBP (a kinase dead version of Pkc1, *pkc1-K853R* (Watanabe et al, 1994)). These plasmids were separately transferred into 159 GFP strains including kinetochore proteins, members of the CWI pathway and actin related proteins (Fig 5 source data). We confirmed that Pkc1-GBP was co-localizing with endogenously GFP-tagged kinetochore using fluorescence imaging. We found that the GFP-GBP binding is sufficient to cause considerable relocalization of Pkc1 to the kinetochore in all cases (Fig 5A), consistent with previous observations (Berry et al, 2016; Olafsson & Thorpe, 2015). We found that a number of haploid strains, which contain only a single endogenously encoded copy of the GFP tagged protein, are restricted, for growth when expressing Pck1-GBP. The strongest SPI caused by Pkc1-GBP is with Bck1-GFP (Fig 5B, Fig 5 source data). Bck1 is the downstream target of Pkc1 in the CWI pathway (Fig 4A). In addition, association of Pkc1 with other MAP kinases further downstream in the CWI pathway also led to a growth defect (Mkk1, Mkk2 and Slt2) (Fig 5B, Fig 5 source data). From the kinetochore, we found that Ndc10, Cse4, Dad3, Dad4, Ctf19, Okp1, Cep3 all gave SPIs with Pck1 (Fig 5B, C and Fig 5 source data). Also, some SPB components such as Spc42 had SPIs with Pkc1 (Fig 5B, Fig 5 source data). To test whether these effects were specific for the kinase activity of Pkc1, we compared the growth defects caused by Pck1-GBP, *pkc1-ha*-GBP and *pkc1-kd-GBP*. We found that downstream members of the CWI pathway are strongly sensitive to either Pck1-GBP or *pkc1-ha*-GBP but not to *pkc1-kd*-GBP, except Bck1-GFP that is sensitive to all three Pkc1-GBP versions (Fig 5C). From the kinetochore proteins, Dad3 showed a strong growth defect with Pck1-GBP or *pkc1-ha*-GBP but not to *pkc1-kd*-GBP (Fig 5C). It is possible that some of these SPIs are caused by mislocalization of the GFP-tagged protein (caused by association with the Pkc1-GBP). To address this we repeated the SPI screen in heterozygously GFP-tagged diploid strains. In these strains there is one allele of a gene linked to GFP and the other allele is not tagged. Thus, if the GFP-tagged protein is mislocalized, then the untagged version may be able to complement the defect. We compared again the growth defects caused by Pck1-GBP, *pkc1-ha*-GBP and *pkc1-kd-GBP* in these heterozygous diploids (Fig 5D and E). We found that the only proteins that produce a SPI when comparing *pkc1-ha*-GBP with *pkc1-kd*-GBP are the four downstream MAP kinases in the CWI pathway: Bck1, Mkk1, Mkk2 and Slt2 (Fig 5D and E). However, when comparing Pkc1-GBP with *pkc1-kd*-GBP, only the immediate downstream MAP kinase (Bck1) is a SPI (Fig 5D and E).

**Figure 5.**
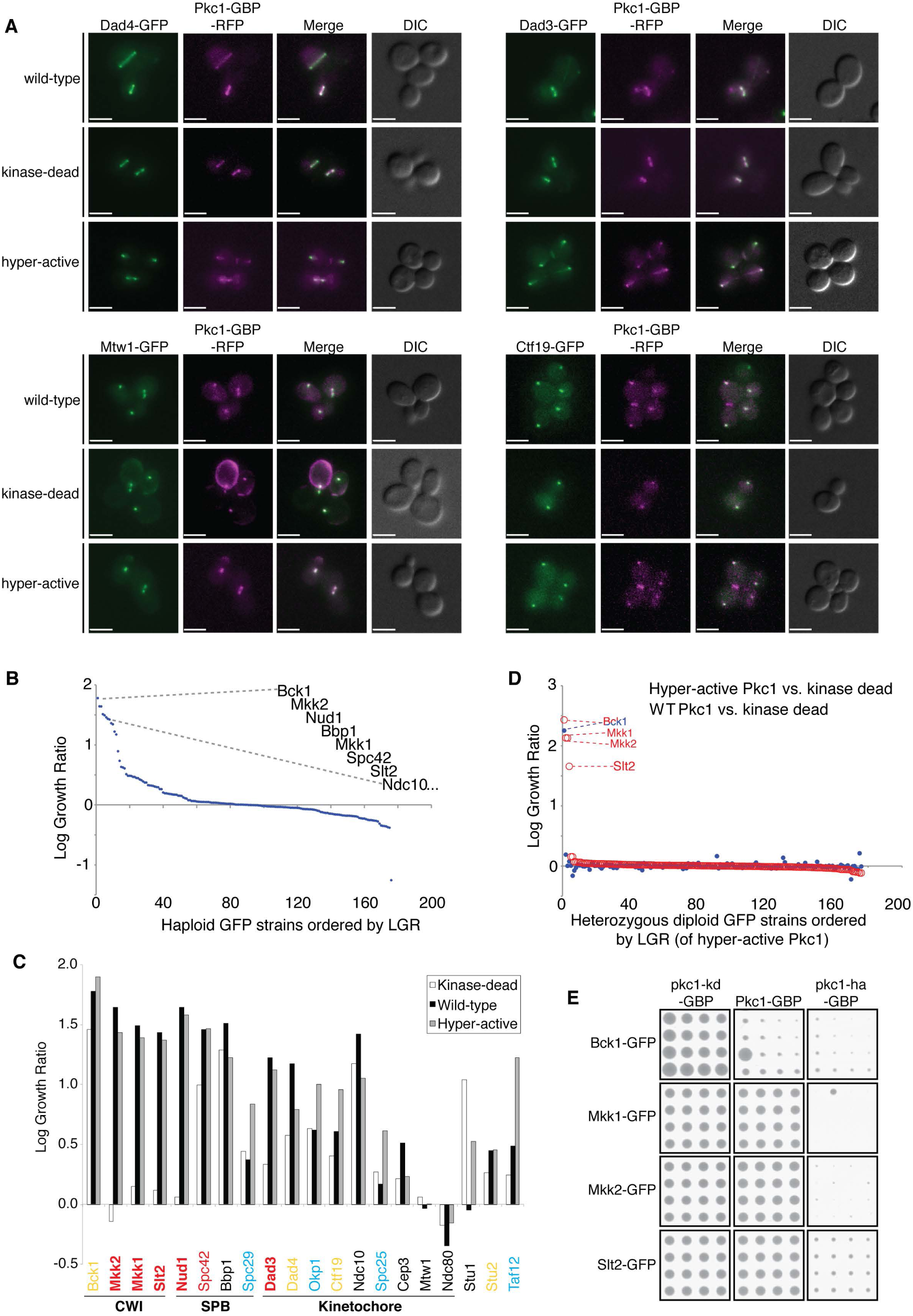
Synthetic Physical Interactions with Pkc1. A The GBP-tagged versions of Pkc1, *pkc1-kd* and *pkc1-ha*, all colocalize with kinetochore proteins as assessed by fluorescence imaging. Four examples are shown, Dad4-GFP, Dad3-GFP, Mtw1-GFP and Ctf19-GFP. B Log growth ratio of GFP tagged strains expressing Pkc1-GBP versus expressing GBP. Pkc1-GBP was transformed into an array of 176 GFP-tagged haploid strains, encoding 159 different GFP proteins (17 strains were present in duplicate). The relative growth of the resulting strains was assessed to detect a mitotic phenotype. A positive ‘Log Growth Ratio’ (LGR) indicates a growth defect. The strongest growth defect was produced by the association of Pkc1 with its kinase target, Bck1 (LGR=1.8). Associations of Pkc1 with several kinetochore proteins also produced growth defects Ndc10 (LGR=1.4), Cse4 (LGR=1.3), Dad3 (LGR=1.2 & 0.9) and Dad4 (LGR=1.2). An LGR of 1 equates with a control colony 2.7 times as large as an experimental colony, LGRs greater than 0.4 are detectable by eye. C Log growth ratio of GFP tagged strains expressing *PKC1-GBP*, *PKC1-KD-GBP* or *PKC1-HA-GBP* versus expressing *GBP* alone. GFP protein associations whose growth effect is specific to both the Pkc1-GBP and *pkc1-ha*-GBP are labeled in red (with bold text indicating a quantitatively stronger relative growth defect). GFP protein associations whose growth effect is specific to *pkc1-ha*-GBP are labeled in blue. GFP protein associations whose growth effect are not specific, but are greater with the Pkc1-GBP and/or *pkc1-ha*-GBP are labeled in yellow. D Log Growth Ratio comparisons of diploid heterozygously GFP-tagged strains expressing either *PKC1-GBP*, *PKC1-KD-GBP* or *PKC1-HA-GBP*. When comparing Pkc1-GBP with *pkc1-kd*-GBP, the only growth defect was in Bck1, the direct target of Pkc1 (blue dots). When comparing *pkc1-ha*-GBP with *pkc1-kd*-GBP, the four kinases in the CWI pathway downstream of Pkc1 Bck1, Mkk1, Mkk2 and Slt2 (red dots) showed a growth defect. E Examples of the strong growth defects produced by combining Pkc1-GBP *pkc1-kd*-GBP or *pkc1-ha*-GBP with GFP-tagged members of the CWI pathway are shown (See Fig 5 source data). Each strain is arrayed in 16 replicates. Source data for this figure is available on the online supplementary information page.

Thus, although we cannot rule out a direct role for Pkc1 at the yeast kinetochore, we find that constitutive recruitment of Pkc1 to kinetochores does not result in a mitotic phenotype consistent with an indirect role of Pkc1. This is further supported by the ability of sorbitol to rescue the *pkc1-1* kinetochore phenotype. These data suggest that *pkc1-1* mutants activate a stress response pathway that indirectly leads to the disassembly of the kinetochore cluster. Since previous work has demonstrated cell stress results in altered MT dynamics (Robertson & Hagan, 2008), we examined whether microtubules were affected by disruption of Pkc1. We used the Pkc1-AID inducible degron system in a strain that contains Dad4-YFP, Spc42-RFP and tubulin (Tub1) tagged at the N-terminus with mTurquois (mTurq-Tub1, a fluorescent version of Tub1 that minimized the perturbation of MT function (Markus et al, 2015)). We induced Pkc1-AID degradation for 3 hours to identify cells as their kinetochore foci start to dissever. We find that a significant proportion of cells have altered kinetochore (Fig 6A) and that these cells have aberrant MTs (Fig 6B). The Pkc1 depleted cells appear to have hyperstable MTs compared with controls, and the extra-kinetochore foci are associated with these MTs (Fig 6B). Thus, these observations support the notion that the kinetochore phenotype seen in Pkc1 deficient cells is associated with altered MT stability. These data therefore provide a link between the stress response pathways and kinetochore function.

**Figure 6.**
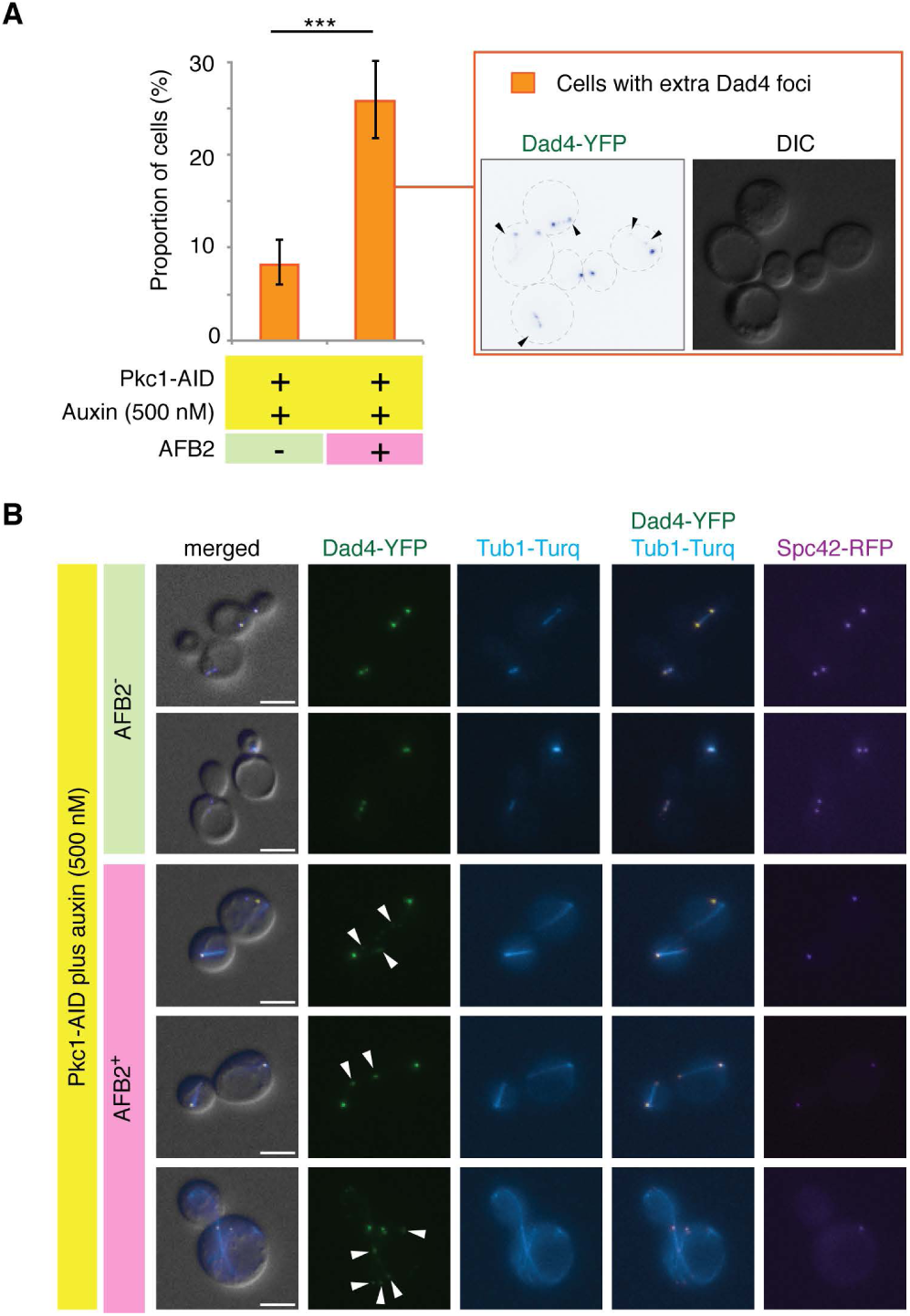
Pkc1 maintains normal microtubule morphology. A Pkc1-AID was depleted in cells by addition of 500 µM auxin. The cells were incubated for three hours to identify cells as their kinetochore foci sub-divide (black arrowheads within the inset panel). The proportion of cells with multiple Dad4-YFP foci is shown; *** indicates a *p* value of 4×10^−14^ using Fishers exact test, and error bars show 95% exact binomial confidence intervals. B Cells that contain additional Dad4-YFP foci (white arrowheads) show colocalization of the Dad4-YFP foci with the microtubules, scale bars are 5 µm.

## Discussion

We have performed a genome-wide screen to determine the pathways regulating the function of the outer kinetochore protein Dad4, part of the DAM1-DASH complex. This complex is an essential component of the outer kinetochore that stabilizes the interaction between kinetochores and the dynamic ends of spindle MTs. The Dad4 screen revealed 315 ORFs that affect the levels and/or localization of this protein at the kinetochore foci. A significant proportion of these mutants also produce a CIN phenotype (Stirling et al, 2011) (enrichment *p*-value = 5×10^−14^) suggesting that they play a role in chromosome segregation. The mutants are also enriched for those that are sensitive to gamma irradiation, hydroxyurea and benomyl (enrichment p-values 2×10^−14^, 9×10^−10^ and 7×10^−8^ respectively). Despite these enrichments, the mutants that perturb Dad4-YFP do not include classical DNA repair genes such as *RAD51* or *MRE11* but rather genes involved in mitosis and checkpoint function (there are some exceptions such as the nuclease gene *APN1*). The mutant list was also enriched for genes involved in RNA metabolism, transport, the nuclear pore and transcription (Fig 1F).

We investigated the role of one particular mutant, *pkc1-1*, in more detail since this evolutionary conserved kinase is important for accurate chromosome segregation (Stirling et al, 2011), yet the Pkc1 kinase has no known role at the yeast kinetochore. We found that loss of Pkc1 led to kinetochore declustering and loss of kinetochore foci (Fig 2 and 3). Recently, a mitotic role for Protein Kinase epsilon (PKCε) has been discovered in mammalian cells (Brownlow et al, 2014; Saurin et al, 2008). PKCε regulates the localization of SAC proteins, BubR1/Mad3 and Bub1 to kinetochores, and in its absence cells progress prematurely through mitosis. These data suggest a direct role for PKCε at the kinetochore. We were not able to detect Pkc1 at the kinetochore using fluorescent imaging (Fig 5SD), nor has Pkc1 been isolated with purified kinetochores (Akiyoshi et al, 2010; Ranjitkar et al, 2010). However, if Pkc1 were to act only transiently or through the action of a small number of molecules, then this would not be detected. To test whether Pkc1 has a direct effect at the kinetochore we used the *Synthetic Physical Interaction* system to forcibly associate Pkc1 to the kinetochore. Although we found that several kinetochore proteins do perturb growth when forcibly bound to Pkc1 (Fig 5C), these are likely due to disruption of the function of essential kinetochore proteins, since the phenotype is not present in heterozygously-tagged diploid cells, which include a non-bound form of the kinetochore protein (Fig 5 source data). However, we cannot rule out a direct role for Pkc1 mediated phosphorylation at the kinetochore, perhaps restricted to a specific point in the cell cycle.

Since the kinetochore phenotype of *pkc1-1* cells was rescued by the addition of the osmo-protectant sorbitol, we suggest that cells in which Pkc1 is perturbed are activating stress response pathways that indirectly mediate the kinetochore phenotype. There is precedent for the stress response pathways altering MT dynamics (Robertson & Hagan, 2008) in fission yeast. The effect of osmotic stress was seen on filamentous actin and microtubules. Consistent with this, we find that MTs appear hyper-stabilized in budding yeast cells depleted of Pkc1 (Fig 6). Thus, these data provide a rationale for the chromosomal instability phenotype seen in *PKC1* mutants. More broadly it may be anticipated that stress response pathways could exert an effect on chromosome transmission via their action on the actin cytoskeleton and MTs. It will be interesting to decipher which stress response pathways contribute to the kinetochore phenotype and to determine whether kinetochore structure is controlled in part by microtubule dynamics. We see hyperactivation of the canonical Hog1 pathway, but since this is active in *pkc1-1* cells at the permissive temperature, there must be other contributing factors.

## Materials and Methods

### Yeast Strains and Growth

A full list of the yeast strains used in this study is provided in Table S1 and plasmids in Table S2. Yeast were manipulated using standard methods (Sherman, 2002); ts strains were grown at 23°C and then at non-permissive temperatures as indicated in the text.

### Synthetic Genetic Array (SGA)

Strains encoding fluorescently tagged spindle components Dad4-YFP and Spc42-RFP (T34 and T37) were mated with an array of gene deletions (Winzeler et al, 1999) or ts alleles (Li et al, 2011) by combining the cells together on YPD plates using a Singer ROTOR pinning robot (Singer Instruments, UK). Diploids were then selected by two cycles of growth on rich media containing geneticin (G418), hygromycin (HYG) and nourseothricin (NAT). These diploids were sporulated for 2 weeks at 23°C on sporulation medium. The resulting spores were selected in four successive cycles of growth for 1 or 2 days at 30°C on synthetic media lacking leucine and lysine (1) with thialysine; (2) with thialysine and G418; (3) thialysine, G418 and NAT; and (4) thialysine, G418, NAT and HYG.

### Fluorescence imaging

Cells were prepared for imaging by growth in liquid synthetic medium containing 120 mg/litre adenine at 23°C for ~5 hours in 96 well plates. The cells were then collected by centrifugation and placed onto an 8×6 grid of agar pedestals (Werner et al, 2009). These 48 pad arrays were covered with a custom-sized 175 µm thick coverslip and imaged on an upright epifluoresence microscope (Zeiss Axiovision Z2), using a 63x 1.4NA plan apochromat oil immersion lens. Zeiss Immersol 518F immersion oil was used with a refractive index of 1.5181. Fluorescence illumination for the screen was provided by a 100W mercury bulb. Fluorescence filter cubes were as follows: YFP imaging used Zeiss filter set 46 (excitation BP 500/20; dichroic FT 515, emission BP 535/30), RFP imaging used Zeiss filter set 63HE (excitation BP 572/25; dichroic FT 590, emission BP 629/62), GFP imaging used Zeiss filter set 38HE (excitation BP 470/40; dichroic FT 495, emission BP 525/50) and CFP imaging used Zeiss filter set 47HE (excitation BP 436/25; dichroic FT455, emission BP 480/40). Later experiments employed a Zeiss Colibri LED illumination system (445 nm for CFP excitation, 470 nm for GFP excitation, 505 nm for YFP excitation and 590nm for RFP excitation), with the filter cubes outlined above. A z stack of 21 images with a vertical separation of 350 nm was acquired for each strain. Brightfield contrast was enhanced with differential interference contrast (DIC) prisms. The resulting light was captured using a Hamamatsu ORCA ERII CCD camera with 6.45 µm pixels, binned 2×2 or a Flash 4.0 Lte CMOS camera with 6.5 µm pixels, binned 2×2 (Hamamatsu Photonics, Japan). The exposure times were set to ensure that pixels were not saturated, typically < 200 msec – for the genome-wide screen Dad4-YFP exposure times were 40 msec and Spc42-RFP 50 msec. Images shown in the figures were prepared using Volocity imaging software (Perkin Elmer Inc., USA).

### Image analysis

Qualitative image analysis was performed using Volocity to visualize the fluorescence images. Quantitative image analysis used an automated ImageJ script, FociQuant to quantify the kinetochore fluorescence signal (Ledesma-Fernandez & Thorpe, 2015). In brief, each kinetochore is identified automatically using the ‘FindMaxima’ function in ImageJ. The fluoresecence intensity within a ~600 nm square box in the focal plane of each foci is measured and a local background signal for each focus is subtracted. We found that the deviations in fluorescence intensity of control strains across each 48 pad imaging plate and consequently we used a smoothing algorithm to normalize values across each plate of the screen (Olafsson & Thorpe, 2015).

### Synthetic Physical Interactions

We used the established method for synthetic physical interactions (Olafsson & Thorpe, 2015). In brief, The *GBP*/*PKC1-GBP* alleles were expressed from a single copy *CEN* plasmid under the control of a *CUP1* promoter, all strains were grown at 30°C. Plasmids encoding the GBP-tagged proteins and controls were separately transferred into the GFP strains using ‘selective ploidy ablation’ (SPA) (Reid et al, 2011). Arrays of 96 strains from the GFP collection were grown on rectangular plates containing YPD media, typically at 1536 colonies per plate density (16 replicates of each strain). After growth overnight, these *MAT***a** plates were copied onto new YPD plates with a lawn of donor *MATα* yeast strain containing a specific *LEU2* plasmid. The resulting diploids were grown overnight on YPD to allow mating. To select for haploid cells, these cells were then copied onto media containing galactose and lacking leucine (GAL – leu) to compromise the conditional chromosomes of the donor strain, while selecting for the specific plasmid. 24 hours later the colonies were replicated to GAL –leu medium containing 5-fluoroorotic acid (GAL –leu 5FOA) to select against the remaining donor chromosomes. To select for diploids, after the YPD mating step, the colonies were copied onto media lacking histidine and leucine (-His, -Leu) to select for both the plasmid and the GFP in the recipient strain. After 24 hours the cells were copied onto fresh –His, -Leu plates. For both haploids and diploids the plates were imaged using a desktop scanner in transmission mode (minimum resolution 300 dpi) after 24-72 hours. Quantitative analysis of the resulting colony size was performed as previously reported. First, the colony sizes on each plate are normalized to a median value of 1 to account for plate to plate variations in growth. Next, for each strain, the average colony size of the control was compared with that of the experimental plasmid(s) to generate a growth ratio.

The natural log of the growth ratios is reported here for each strain; LGR = ln (control size/experimental size). An LGR of 0 indicates equal size colonies on experiment and control, whereas an LGR of 0.4 equates with an experimental colony 67% of the size of a control colony.

### Protein extraction and Western Blot Analysis

Whole cell extracts where prepared by using trichloracetic acid (TCA) as previously described (Olafsson & Thorpe, 2015). Protein extracts were separated on acrylamide gels (7.5% for Pkc1 and 10% for Dad3, Dad4 and Mtw1). Then, gels were transferred to 0.45 μm supported nitrocellulose membrane (BIORAD). Membrane blocking and antibody incubation were performed using Western Blocking reagent (Roche) as recommended by manufacturer. Anti-FLAG antibody (Sigma, F7425) and anti-GFP antibody (Roche, 11814460001) were used at 1:1000 dilutions. Anti-Pgk1 antibody (Invitrogen, 459250) was used at 1:10000.

HRP conjugated anti-Rabbit IgG (Sigma, A0545) and anti-mouse IgG (Abcam, ab97265) were used at 1:100000 and 1:30000 dilution, respectively. Membranes were incubated for 1 min with Lumi-Light Western Blotting substrate (Roche) before film exposure.

## Acknowledgements

We would like to thank Brenda Andrews, Lisa Berry, Erika Aquino, Helle Uhlrich, Wei-Lih Lee, Iain Hagan, Peter Parker and Zemmer Gitai for materials, assistance and critical advice on this manuscript. This work was largely funded by the United Kingdom Medical Research Council (MRC) MC_UP_A252_1027. This work was also supported by the Francis Crick Institute which receives its core funding from Cancer Research UK (FC001183), the UK Medical Research Council (FC001183), and the Wellcome Trust (FC001183). Our images are hosted by the Image Data Repository, which is funded by the BBSRC (BB/M018423/1).

